# Information and communication technology, mobile devices, and medical education

**DOI:** 10.1101/420281

**Authors:** Andrea Rodríguez-Ríos, Gerardo Espinoza-Téllez, José Darío Martínez-Ezquerro, Mario Enrique Rendón-Macías

## Abstract

**Introduction:** Information and communication technologies (ICT) are practical and highly available tools. In medical education, ICTs allow physicians to update their knowledge and remember the necessary information within reach of current mobile devices. ICTs as preparation tools for medical education have not been reported for medical students in Mexico.

**Methodology:** To assess the use of mobile devices as ICTs with medical education purposes, we distributed a questionnaire through an online survey management system to all the medical students (n=180) from a private university in Mexico City, 100% agreed to participate. We developed a questionnaire based on previous surveys and adapted it to our university.

**Results:** All participants reported possession of an electronic mobile device, and 95% used it regularly for learning purposes. Regardless of the school year, the most frequent usage given to these devices was the search and reading of medical articles, the use of medical calculators, and taking notes. As the levels in career advances, there was a reduction in the use of electronic devices. According to the students, the main barriers towards using mobile devices for learning purposes were both the lack of access to the Internet and permission from the professor to use them.

**Conclusion:** Most medical students use mobile devices for learning purposes, but usage changes during their education. It is convenient to encourage the use of mobile devices and the development of ICT skills as tools for educational purposes rather than banning their use in schools and hospitals.

## Introduction

The high-speed information and knowledge development of our current society makes the use of electronic resources and devices an essential skill for the medical profession (1) for both learning and updating medical knowledge (2–5). In the last two decades, medical students have to consult thousands of scientific articles as sources of new knowledge (6,7). As a consequence, there is a growing interest in information and communication technologies (ICT) skills during student’s basic and clinical training (8). In this regard, Samuel and colleagues in Tanzania (6) reported a perception of inferior performance compared to other students of medical schools from developed countries; the 92 students surveyed attributed that perception to the lack of ICT, mainly personal computers. A similar study in Malaysia (9), where the students had reported that 53% of their students claimed to use them for educational purposes with varying times, half of the students used computers between 1-2 hours, 6.5% between 3-6 hours, and 3.4% more than 6 hours per day. However, none of the authors explored the use of these technologies as a means of accessing medical information. Other authors such as Hye Won Jang and Kyong-Jee Kim (10) have assessed the use of videos as an ICT learning strategy; the authors evaluated the impact of a series of videos designed to respond favorably to the Objective Structured Clinical Examination (OSCE) among students from 34 Korean universities. After watching videos, they answered an exam and requested their opinion on its usefulness to solve the examination. Almost all the students who answered the survey (91.9%) agreed with the usefulness of the videos. It is relevant to mention that one-third of the participants watched the videos on mobile devices and considered it the most accessible way to do so. However, they also pointed out the difficulty of interacting with these devices and software. More recently, improved applications for clinical practice of both physicians and medical students have been developed for computers as well as mobile devices.

In 2014, Boruff and Storie (11) evaluated the appropriateness of Canadian libraries to facilitate the interaction between the members of their universities, students and teachers, with the availability and accessibility of information collections. In general, they found that doctors, both in training and teachers, had no problem accessing collections but ignored the most reliable sources of information and had limited access to quality or authenticated databases.

Despite the high frequency of access to medical information and communication technologies, there are no studies in Mexico regarding the use of these tools by medical students.

In contrast to other studies that focused on basic applications of computers or on the distraction generated by this type of technology (12–15), the aim of our study was to analyze the specific use of information and communication technologies as learning tools in medical education, eliminating factors such as its use for entertainment or distraction, or internet accessibility (Wifi) in school or at the hospital.

## Materials and methods

An observational, cross-sectional, descriptive, and prospective study was carried out during the period of January to March 2016. We invited medical students from our private medical school to participate in the study. Since all students had mobile devices, we created a survey using an online platform (https://www.surveymonkey.com/). Students enrolled in the sixth year of medical school were not eligible for this study as they move to rural areas to complete their social service assignments, where access to the Internet is limited or absent.

Questionnaire construction. The questionnaire consisted of 36 Open-ended and Multiple-choice questions divided into three sections: general information, information technology in the school, and information technology in the hospitals (4, 22, and 10 questions, respectively) (Online Resource 1; available at https://osf.io/tnx4d/). Finally, the mandatory thirty-sixth question referred to the informed consent for the research use of the information provided by each participant. Some questions allowed more than one answer. The questionnaire was constructed for our purposes, considering some questions from similar surveys conducted in other countries (11,15–17). Three professors not involved in the project reviewed the clarity and content of the questionnaire. We sent the questionnaire through an online survey management system using the institutional e-mails from all the students enrolled in the school year of 2016 (n= 180). From the invitation date, the page remained open for two months and only accepted one response per device; participants were able to make changes at any time. To encourage participation, we informed on the importance of the study and reported the response rate of each school grade for motivation. Statistical analysis. The information was collected directly to a database available in the same software and transferred to a commercial database for further analysis. At all times, the anonymity of the participants was protected by requesting only information on age, gender, and scholar year. We ensured one-time participation by restricting access by the device IP (internet protocol) address. For each of the qualitative variables, the findings were summarized in frequencies and percentages. To maintain anonymity, we analyze the data by scholar year and place of academic activities. The mean and its standard deviation were obtained for the qualitative variables. To identify statistical differences in means we used a parametric test (Student’s t-test), and chi-square test for proportions, considering significant differences with a p-value less than 0.05.

## Results

All eligible medical students enrolled in 2016 (n=180) participated in the online survey. The distribution of participants, according to the school year (Table 1). We had a higher number of students in the second and fourth years and lower numbers in the first and fifth years of medical school. The mean age for both men and women was 21 years old, with no differences between school years.

**Table 1.** Demographic characteristics of students participating in the online survey (n=180)

| Variable | Males<br>n=88 | Females<br>n=92 | All<br>n=180 |
| --- | --- | --- | --- |
| **School year\*** |  |  |  |
| 1st | 14 (51.9%) | 13 (48.1%) | 27 (9%) |
| 2nd | 19 (42.2%) | 26 (57.8%) | 45 (25%) |
| 3rd | 21 (46.7%) | 24 (53.3%) | 45 (25%) |
| 4th | 24 (54.5%) | 20 (45.5%) | 44 (24.5%) |
| 5th | 10 (56.6%) | 9 (47.4%) | 19 (10.5%) |
| **Age in years\*\*** |  |  |  |
| Mean ± 1 SD | 21 ± 1.7 | 21 ± 1.8 | 21 ± 1.8 |

\*Pearson's chi-square test 1.6, df=4, p= 0.79; \*\*Student t test -0.28, df=178, p=0.77

### Use of electronic devices for information and communication technology

All students reported owning at least one type of electronic device. All of them had a smartphone, 82.8% (149/180) a laptop, and 98% an electronic tablet.

### Use of information technology

95% of medical students reported using their devices for learning purposes (improving comprehension, memory, and school performance), while only 5% did not use their devices for those purposes. Due to the observed similarity by gender (p = 0.77; Table 1), the use of ICTs was compared only by the school year and according to the places where they attended classes.

Table 2 shows the use of electronic devices on campus, according to the school year. Our data revealed that the primary use of laptops and tablets was taking notes, especially in the first four school years, followed by tablets or phones for reading medical articles or e-books (> 80% of students), regardless of the school year. The search for information on drugs and medical update platforms increased from the second school year. Inquiry of medical practice guidelines was more common among third- and fourth-year medical students (clinical courses), followed by searches for differential diagnoses. Likewise, there was an increase in the use of medical calculator applications for clinical conditions, which were rarely used in the first two years, but not in the last three years. Finally, about 50% of the students reported using ICTs as a means of communication within the university campus. The use of electronic devices in hospitals was also analyzed and revealed that the primary purpose was to search for medical online calculation platforms (Table 3). This use increased significantly over the scholar years. Also, they used ICTs for reading articles or books, mainly in the last two scholarly years. Taking notes with electronic devices increased in the third and fourth years, and sharply reduced in the fifth year (medical internship). The search for drug information, although remaining high (> 66%), was no longer as high as in the time spent in the university campus (> 90%). Searches for clinical practice guidelines were higher in advanced school years, especially when internet access was available at the hospitals. The communication’s use of electronic devices remained stable over the years and in equal proportion as when they were in or outside the university campus.

**Table 2.** Use of electronic devices in the university campus by school year.

| Laptop, computer, smartphone or tablet were used for... | Medical school year |  |  |  |  | p-value** |  |  |
| --- | --- | --- | --- | --- | --- | --- | --- | --- |
|  | 1st<br>n= 15 | 2nd<br>n= 38 | 3rd<br>n= 39 | 4th<br>n= 42 | 5th<br>n= 11 | All<br>n=145 | X<sup>2</sup>-P | X<sup>2</sup>-T |
| taking notes | 86.7% | 94.7% | 94.9% | 95.2% | 54.5% | 90.3% | **0.0004** | 0.10 |
| reading articles or medical books | 86.7% | 97.4% | 97.4% | 95.2% | 81.8% | 93.1% | 0.16 | 0.69 |
| drugs information search | 66.7% | 94.7% | 97.4% | 97.6% | 90.9% | 92.4% | **0.0008** | **0.01** |
| search for information platforms | 53.3% | 92.1% | 92.3% | 92.9% | 90.9% | 87.5% | **0.006** | **0.008** |
| search clinical practice guidelines | 46.7% | 52.6% | 74.4% | 88.1% | 54.5% | 65.5% | **0.002** | **0.005** |
| communication | 46.7% | 52.6% | 56.4% | 50% | 45.5% | 51.0% | 0.94 | 0.89 |
| medical calculations | 6.7% | 42.1% | 79.5% | 78.6% | 81.8% | 62.0% | **0.0017** | **0.005** |
| differential diagnoses search | 0% | 44.7% | 35.9% | 57.1% | 9.1% | 37.2% | **0.0001** | 0.31 |

\*\* X<sup>2</sup>-P, Pearson's chi-square test; X<sup>2</sup>-T, Trend chi-square test

**Table 3.** Use of electronic devices in hospitals by school year*.

| Laptop, computer, smartphone or tablet were used for... | Medical school year |  |  |  |  | p-value** |  |
| --- | --- | --- | --- | --- | --- | --- | --- |
|  | 2nd<br>n= 21 | 3rd<br>n= 37 | 4th<br>n= 35 | 5th<br>n= 9 | All<br>n=102 | X<sup>2</sup>-P | X<sup>2</sup>-T |
| search for information platforms | 76.2% | 73% | 88.6% | 88.9% | 80.4% | **0.04** | 0.09 |
| reading articles or medical books | 76.2% | 81.1% | 74.3% | 88.9% | 78.4% | 0.76 | 0.80 |
| taking of notes | 66.7% | 89.2% | 82.9% | 44.4% | 78.4% | **0.01** | 0.57 |
| drugs information search | 66.7% | 86.5% | 88.6% | 88.9% | 83.3% | 0.24 | 0.14 |
| medical calculations | 57.1% | 81.1% | 82.9% | 66.7% | 75.5% | 0.11 | 0.21 |
| search clinical practice guidelines | 52.4% | 70.3% | 85.7% | 88.9% | 73.5% | **0.03** | **0.004** |
| differential diagnoses search | 47.6% | 35.1% | 51.4% | 11.1% | 41.2% | 0.12 | 0.47 |
| communication | 42.9% | 48.6% | 37.1% | 55.6% | 50.0% | 0.59 | 0.92 |

\*This question allowed more than one answer
\*\* X<sup>2</sup>-P, Pearson's chi-square test; X<sup>2</sup>-T, Trend chi-square test
\*\*\*The reported use of electronic devices for study purposes in the hospital was almost 60% (102/180)

### Duration of use of electronic devices for educational purposes

Table 4 summarizes the proportion of students who invested less than 24 hours, from 24 to 72 hours, and more than 72 hours per week in learning with the support of their electronic devices. Except for third-year students, they usually accessed their devices for less than 24 hours per week, especially during the fifth year. In their third school year, just over half of the students used their devices between 24 to 72 hours. A duration above 72 hours was observed in fourth-year students, although the difference was not statistically significant.

**Table 4.**
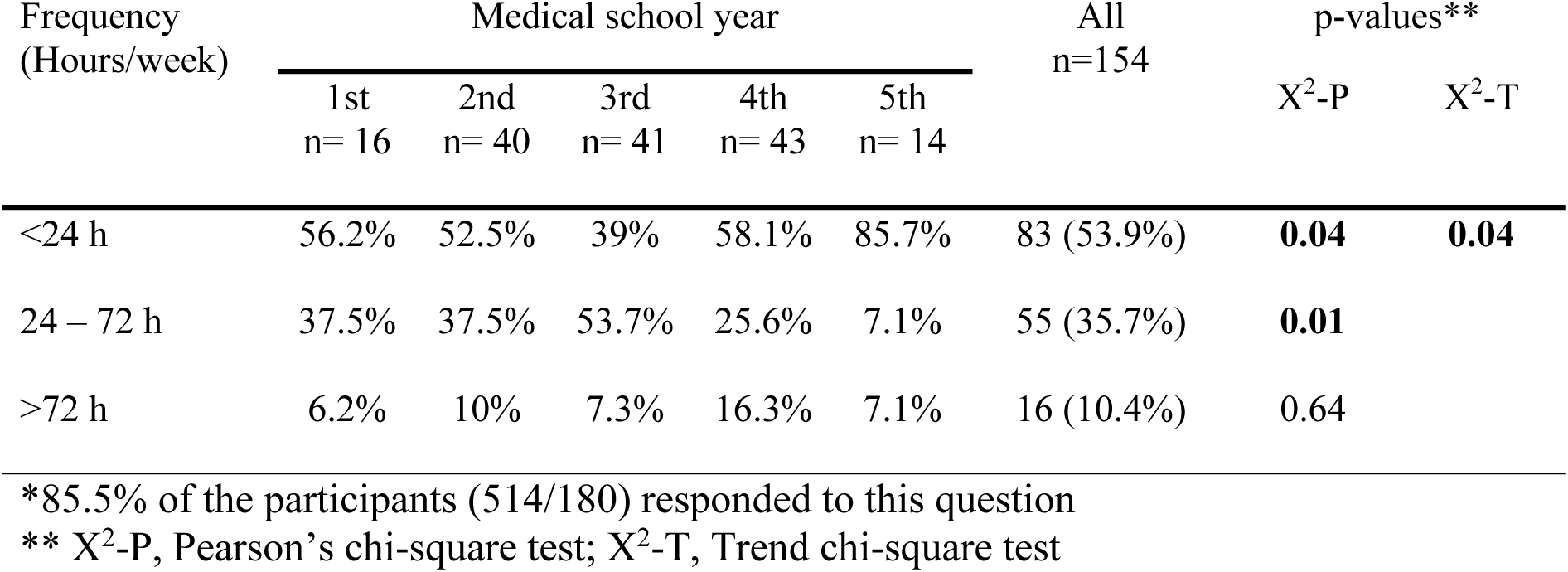
Time spent using electronic devices for learning purposes.

| Frequency<br>(Hours/week) | Medical school year |  |  |  |  | All<br>n=154 | p-values** |  |
| --- | --- | --- | --- | --- | --- | --- | --- | --- |
|  | 1st<br>n= 16 | 2nd<br>n= 40 | 3rd<br>n= 41 | 4th<br>n= 43 | 5th<br>n= 14 |  | X<sup>2</sup>-P | X<sup>2</sup>-T |
| <24 h | 56.2% | 52.5% | 39% | 58.1% | 85.7% | 83 (53.9%) | **0.04** | **0.04** |
| 24 – 72 h | 37.5% | 37.5% | 53.7% | 25.6% | 7.1% | 55 (35.7%) | **0.01** |  |
| >72 h | 6.2% | 10% | 7.3% | 16.3% | 7.1% | 16 (10.4%) | 0.64 |  |

\*85.5% of the participants (514/180) responded to this question
\*\* X<sup>2</sup>-P, Pearson's chi-square test; X<sup>2</sup>-T, Trend chi-square test

### Barriers to the use of electronic devices

The barriers were different for the university campus and the hospital headquarters (Table 5). Students mentioned that the main obstacles in the university campus were the instability of the internet connection, followed by the lack of authorization from the professors, being in areas where there was no network signal, or did not have access (password permission), as well as the lack of time for their consultation. All of the students (180/180) referred to difficulties in the use of electronic devices at the university campus. On the other hand, the limitations of using electronic devices in hospitals were the absence of internet access or the refusal of the authorities to provide the access codes and the non-authorization by the professors. Other less mentioned restrictions were that they were unaware of the resources or how to access them, the little time available, and connection instability. The least common limitation was the lack of information on how to use some platforms provided by the hospitals. Overall, 85% of the students (153/180) referred to difficulties in the use of electronic devices at the hospitals.

**Table 5.** Obstacles to access electronic information sources in medical learning settings.

| Barriers | University campus<br>n= 180* |  | Hospitals<br>n= 153** |  | *p-value\*\** |
| --- | --- | --- | --- | --- | --- |
|  | N | % | N | % |  |
| Technical problems with the connection | 63 | 35% | 23 | 15% | **<0.001** |
| No network or access key | 43 | 23.9% | 108 | 70.6% | **<0.001** |
| Lack of knowledge on the use of<br>resources | 43 | 23.9% | 8 | 5.2% | **<0.001** |
| Lack of time | 20 | 11.1% | 36 | 23.5% | **0.004** |
| Unawareness of available resources | 42 | 23.3% | 34 | 22.2% | 0.84 |
| Not authorized use by the professor | 47 | 26.1% | 42 | 27.5% | 0.87 |

\*\* Pearson's chi-square test with Yates's continuity correction

## Discussion

The data obtained here supports the notion of an extensive use of electronic devices in medical learning settings by students along the school years. Also, we observed a transition in its use from the basic sciences to the clinical courses. In the former, the students use ICTs for pharmacological information searches while in the clinical courses, for clinical practice guidelines searches, particularly in the fifth year of clinical courses where students perform acting internship, the use of ICTs was focused in medical calculators (resolution of formulas of indicators such as “glomerular filtration” or “calculation of body surface”, among others). Taking notes with the support of electronic devices remained an everyday activity throughout the medical school.

One observation that caught our attention was the decrease of searches for clinical practice guidelines in more advanced students (fifth-year acting interns). A possible explanation could be the increased experience and knowledge acquired during the previous school years. A previous study (11) also reported that drug information searches, the use of medical calculators, and taking notes were among the main activities and reasons for using electronic devices. Besides, the authors observed a reduction in the consultation of these digital sources by students and residents, which could be associated with the generational learning gap of these technological devices. Although we did not evaluate the use of electronic devices by professors, we could also expect that these professors do not consult these digital sources as well, given the student-reported barrier (carried-out by the professors) to use these devices. However, future professors may reverse this situation as former medical students and active users of ICTs. Even though this was an exploratory study, medical students reported that the lack of authorization from teachers was a barrier to the exploitation of ICTs. The students referred that this limitation usually occurs during the direct attention of the patients, probably and appropriately to avoid the perception of a distant communication from students towards patients; on some occasions, it also occurred during class.

Extensive use of electronic devices is promoted as useful tools for both clinical learning activities and patient care support, especially in developed countries (16,17). The main goal for the use of ICTs is the immediate and accurate information with the aim of solving problems and doubts during the treatment of a patient. As a result, there has been a development of several applications or “apps” available for smartphones, tablets, or laptops to support information query and decision making by physicians. However, barriers to take advantage of these ICTs may come from the underutilization while teaching and performing clinical practice. In this regard, Boruff y Storie (11) reported that although the main impediment for accessing information is the absence of internet access in clinics and hospitals, ignorance of resources was the second barrier for its use. These authors also found that the lack of training could lead to unreliable information; therefore, they recommend access only to approved medical information. To address this issue, we consider that directed incorporation of ICTs supported by both institutions and professors would improve the usability of these resources as medical learning tools, the quality of patient care, and even counteract the ludic use of electronic devices, at least in medical settings. And again, even when this was an exploratory study, medical students reported that the lack of authorization from professors was a barrier to the exploitation of ICTs.

This could be the result of a negative perception of the use of electronic devices; instead of considering them as useful resources for learning and medical purposes. On many occasions, when the search for information is necessary and beneficial, its utility can overlap the lack of support or permission (17).

In this study, we also found that the use of electronic devices was more frequent in the university campus, probably as a result of internet access availability, in contrast to the clinical or hospital areas, where students experienced difficulties in accessing the Internet or using their devices. However, connectivity failures at the university campus represent a technical barrier when network saturation occurs due to high demand. In addition, the lack of time was a personal barrier sometimes related as a reason for not using ICTs, even when the resource was available. Time was also reported previously as a substantial factor for not accessing electronic information sources (11). Time is even shorter in clinical settings, where the usability could increase during patient rounds accompanied by evidence-based medicine questions for the patient’s care. Their professors could supervise its use in the consultation of drug doses, review calculations of scales or laboratory results, calculations of diagnostic probabilities, among others. In addition to the absence of internet connection in several clinical settings, other common barriers reported by the students include the refusal to provide the internet access codes as well as the risk of thefts. The implementation of computer rooms in clinical settings, as is the case in medical libraries from developed countries, could avoid these barriers (7).

An important aspect of this survey is the recognition that information about new educational techniques based on the interaction of students with digital information sources is still scarce. Problem-based learning methods, as well as simulators or algorithms, through apps or programs, would allow both professors and students, to achieve meaningful learning of the medical topics. A recent study found great interest among students and professors to incorporate these tools for learning and teaching, respectively; however, its implementation was affected by the age and occupation of the professors, the scarce training in their use in addition to the institutional policy about its incorporation (8).

The aim of this study was to explore the use of information and communication technologies during the training of medical students. We consider among its main strengths both the participation of all eligible medical students (100%) as well as its reliability since the information was anonymous and without repercussion in the qualification of the participants. However, we accept several limitations. The barriers reported by our students for the use of ICTs may not occur in other universities and clinical settings. We did not design the questionnaire to explore differences in the use of ICTs for learning purposes categorized by type of device (cell phone, tablet or laptop). We do not know if there are already medical programs in our country designed for the use of electronic devices and ICTs in teaching-learning processes. More studies are necessary for a detailed analysis, in particular, to explore the reasons from the professors to prohibit the use of ICTs, which may be detrimental to the student’s performance if used for distracting non-learning purposes (18), and if it is context-specific. For the moment, we believe that training in their use may improve the learning process of future physicians in the benefit of patient care.

## Conclusions

Most medical students surveyed from a private medical school located in Mexico City own at least one electronic device and use them for learning purposes in both the university and clinical settings. During their career, the use of ICTs transit from theoretical information queries to the search for medical guides and articles, and finally, to medical calculators. The main reported barriers to its use included accessibility to an internet connection (real or imposed) and time availability. The incorporation of these devices and medical applications as teaching-learning tools will depend on the consent of professors and the support from institutional programs and medical settings.

## Declarations

### Availability of the data and material

The questionnaire is available as a supplementary file (Online Resource 1) at the OSF platform https://osf.io/tnx4d/.

### Ethics approval

All procedures performed in this study were under the ethical standards of the international and national research committee and with the 1964 Helsinki declaration and its later amendments or comparable ethical standards. The project received approval by the research and ethics committee of the medical school.

### Consent to participate

All students signed informed consent to participate in the study.

### Authors’ contributions

MERM was responsible for the conceptualization, design of methodology, and supervision; ARR and GET carried out the investigation; ARR, GET, JDME, and MERM prepared the writing of the initial draft; JDME and MERM performed the formal analysis, reviewed and edited the final manuscript. All authors read, approved the final manuscript, gave explicit consent to submit, and agreed to be accountable for all aspects of the work in ensuring that questions related to the accuracy or integrity of any part of the work are appropriately investigated and resolved.

## References

1. Ward JPT, Gordon J, Field MJ and Lehmann HP: Communication and information technology in medical education. Lancet 357: 792–796, 2001.

2. Romanov K and Aarnio M: A survey of the use of electronic scientific information resources among medical and dental students. BMC Med Educ 6, 2006.

3. Sclafani J, Tirrell TF and Franko OI: Mobile tablet use among academic physicians and trainees. J Med Syst 37: 9903, 2013.

4. Eggermont S, Bloemendaal PM and van Baalen JM: E-learning any time any place anywhere on mobile devices. Perspect Med Educ, 2013.

5. Friederichs H, Marschall B and Weissenstein A: Practicing evidence based medicine at the bedside: a randomized controlled pilot study in undergraduate medical students assessing the practicality of tablets, smartphones, and computers in clinical life. BMC Med Inform Decis Mak 14: 113, 2014.

6. Samuel M, Coombes JC, Miranda JJ, Melvin R, Young EJW and Azarmina P: Assessing computer skills in Tanzanian medical students: an elective experience. BMC Public Health 4: 37, 2004.

7. Bravo LE: From printing to Scielo and Pubmed Central. Colomb Med 45: 5–6, 2014.

8. Fan S, Radford J and Fabian D: A mixed-method research to investigate the adoption of mobile devices and Web2.0 technologies among medical students and educators. BMC Med Inform Decis Mak 16, 2016.

9. Lim TA, Wong WH and Lim KY: Perceived skill and utilisation of information technology in medical education among final year medical students, Universiti Putra Malaysia. Med J Malaysia 60: 432–440, 2005.

10. Jang HW and Kim K-J: Use of online clinical videos for clinical skills training for medical students: benefits and challenges. BMC Med Educ 14: 56, 2014.

11. Boruff JT and Storie D: Mobile devices in medicine: a survey of how medical students, residents, and faculty use smartphones and other mobile devices to find information. J Med Libr Assoc 102: 22–30, 2014.

12. Kir T, Ogur R, Kilic S, Tekbas OF and Hasde M: How Medical Students Use the Computer and Internet at a Turkish Military Medical School. Mil Med 169: 976–979, 2004.

13. Maroof KA, Parashar P and Bansal R: How are our medical students using the computer and internet? A study from a medical college of north India. Niger Med J 53: 89–93, 2012.

14. Ayatollahi A, Ayatollahi J, Ayatollahi F, Ayatollahi R and Shahcheraghi SH: Computer and Internet use among Undergraduate Medical Students in Iran. Pak J Med Sci Q 30: 1054–1058, 2014.

15. Nalliah RP and Allareddy V: Students distracted by electronic devices perform at the same level as those who are focused on the lecture. PeerJ 2: e572, 2014.

16. Cheston CC, Flickinger TE and Chisolm MS: Social media use in medical education: a systematic review. Acad Med 88: 893–901, 2013.

17. Masters K and Al-Rawahi Z: The use of mobile learning by 6th-year medical students in a minimally-supported environment. J Int Assoc Med Sci Educ 3: 92–97, 2012.

18. Carter SP, Greenberg K and Walker MS: The impact of computer usage on academic performance: Evidence from a randomized trial at the United States Military Academy. Econ Educ Rev 56: 118–132, 2017.

